# Fine human genetic map based on UK10K data set

**DOI:** 10.1101/809020

**Authors:** Ziqian Hao, Pengyuan Du, Yi-Hsuan Pan, Haipeng Li

## Abstract

Recombination is a major force that shapes genetic diversity. Determination of recombination rate is important and can theoretically be improved by increasing the sample size. However, it is challenging to estimate recombination rates when the sample size is extraordinarily large because of computational burden. In this study, we used a refined artificial intelligence approach to estimate the recombination rate of the human genome using the UK10K human genomic dataset with 7,562 genomic sequences and its three subsets with 200, 400 and 2,000 genomic sequences under the Out-of-Africa demography model. We not only obtained an accurate human genetic map, but also found that the fluctuation of estimated recombination rate is reduced along the human genome when the sample size is increased. UK10K recombination activity is less concentrated than its subsets. Our results demonstrate how the sample size affects the estimated recombination rate, and analyses of a larger number of genomes result in a more precise estimation of recombination rate.

## Introduction

Recombination is a foundation of evolution^1^. The study of recombination helps to decipher mechanisms of linkage disequilibrium^2–4^, nucleotide diversity^5;6^, population admixture and differentiation^7–10^, natural selection^11–14^, and diseases^15;16^. Therefore, much attention has been paid to the determination of the recombination rates of various regions and identification of recombination hotspots in the human genome. Since pedigree studies^17;18^ and sperm-typing studies^19–21^ are technically challenging, indirect approaches, such as those using population genetic methods^22–26^, are more commonly used. Population genetic methods estimate recombination rates based on variations in genomic DNA sequences of a selected population. The rationale behind these indirect approaches is the assumption that coalescent tree length and the number of recombination events occurred in the sample during evolution can be (indirectly) inferred.

According to the coalescent theory, the expected coalescent tree length and number of recombination events in the sample increases when the number of genomes (sample size) analyzed is increased. It has been suggested that the inference accuracy of recombination rates can be improved by using a larger sample size^27–30^. However, it is technically challenging to estimate recombination rates when the sample size is extraordinarily large because of computational burden. In this study, we refined FastEPRR^27^, an extremely fast artificial-intelligence-based software, to estimate the recombination rates of various regions in the human genome using the UK10K data set (*n* = 7,562 genomic sequences)^31^ and its three subsets with 200, 400 and 2,000 genomic sequences under the Out-of-Africa demographic model^32^. We found the smaller fluctuation of local recombination rate along the human genome and less concentrated recombination activity as the sample size was increased. Furthermore, our findings are robust with the assumption of demographic history since the same conclusion was made under the constant population size model. Therefore, the large sample size improves the accuracy of estimated recombination rate, and a fine human genetic map is obtained.

## Materials and Methods

### Fine-tuning FastEPRR

We have recently developed the software “Fast Estimation of Population Recombination Rates” (FastEPRR) using an artificial intelligence method to estimate recombination rate (ρ) based on single nucleotide polymorphism (SNP) data and the finite-site model^27^. FastEPRR first calculates compact folded site frequency spectrum (SFS)^33^ and four recombination-related statistics for sliding windows. A training data set is then generated using a modified version of Hudson’s *ms* simulator^34^, which generates samples conditional on the compact folded SFS under neutral models. Subsequently, the gamboost package^35^ is used to fit the training set and build the regression model. Recombination rate (ρ) is then inferred by the summary statistics of the real data. FastEPRR has been implemented in the R software package^36^ and can estimate recombination rate under different demographic models.

In this study, the FastEPRR software was fine tuned. To optimize speed, for-loops were replaced by “apply” family of functions because R language is greatly effective in carrying out vectorized operations. Input and output operations were minimized to increase the speed. Another modification was to enable the use of different ρ values to generate training samples, determination of the number of independent training samples and estimation of confidence interval. All of these improvements and updates have been incorporated in FastEPRR 2.0.

### Validation of FastEPRR 2.0 with a very large sample size

The neutral data was simulated by using Hudson’s *ms* simulator^34^, a software widely used in the studies of humans and other species. It generates simulated polymorphic data set conditional on mutation and recombination rate under neutral demographic scenarios, according to Kingman’s coalescent process^37^. The following parameters were used: *θ* (population mutation rate) = 4*Nμ* = 30; *ρ* (population recombination rate) = 4*Nr* = 10, 20, 30, 40, 50, 60, 70, 80, 90, 100, 110, 120, 130, 140, and 150; and *n* = 8,000; where *N* is effective population size, *μ* is mutation rate per chromosomal region per generation, and *r* is recombination rate per chromosomal region per generation. For each *ρ*, the mean and the standard deviation were determined using 100 simulated data with specific *ρ* and *θ*.

To determine the accuracy in the estimation of recombination rates on potential recombination hotspot regions, the simulated data were established using a set of large *ρ* values (i.e., 500, 1000, 2000, 3000, 4000, and 5000). The mean and the standard deviation were determined using 10 simulated data for each *ρ* value.

### Building the UK10K genetic map

A total of 7,562 genomes (from 3,781 unrelated individuals) in the UK10K data set were analyzed. To build genetic maps, the window size was set as 10 kb, and the step length was 5 kb. Indels were removed. Windows were excluded if they overlapped with known sequencing gaps in the human reference genome (hg19) or the number of non-singleton polymorphic sites was smaller than 10. The default value set of ρ (0, 0.5, 1, 2, 5, 10, 20, 40, 70, 110, 170, 180, 190, 200, 220, 250, 300, and 350) was used to simulate the training sets.

To estimate recombination rates, the Out-of-Africa model was used^32^. To evaluate the effect of demographic history, the constant size model was also considered. The linear extrapolation was employed if the estimated value is beyond the training set boundary. To ensure the accuracy of results, the default training set values of ρ were adjusted, and FastEPRR was re-ran until the estimated values fell into the range of the given set of ρ. The previous method in FastEPRR was used to evaluate the effects for variable recombination rates within windows^28^. Three data sets with 200 (UK10K-sub200), 400 (UK10K-sub400) and 2,000 (UK10K-sub2000) genomes randomly chosen from the UK10K data set were also analyzed.

## Results

FastEPRR has been shown to perform well with different sample sizes (*n* = 50, 100, 200 and 1,000 chromosomes), and the standard deviation becomes smaller when the sample size is increased^27^. However, its effectiveness on the analysis of a large sample size (e.g., *n* = 7,562 chromosomes) has not been determined. Therefore, the accuracy of recombination rate estimated by FastEPRR was examined when the sample size is extremely large (Figure 1A and 1B). The FastEPRR estimate is unbiased, and its standard deviation becomes much smaller. Thus the inference accuracy of recombination rate is dramatically improved when a greatly large sample size is available.

**Figure 1.**
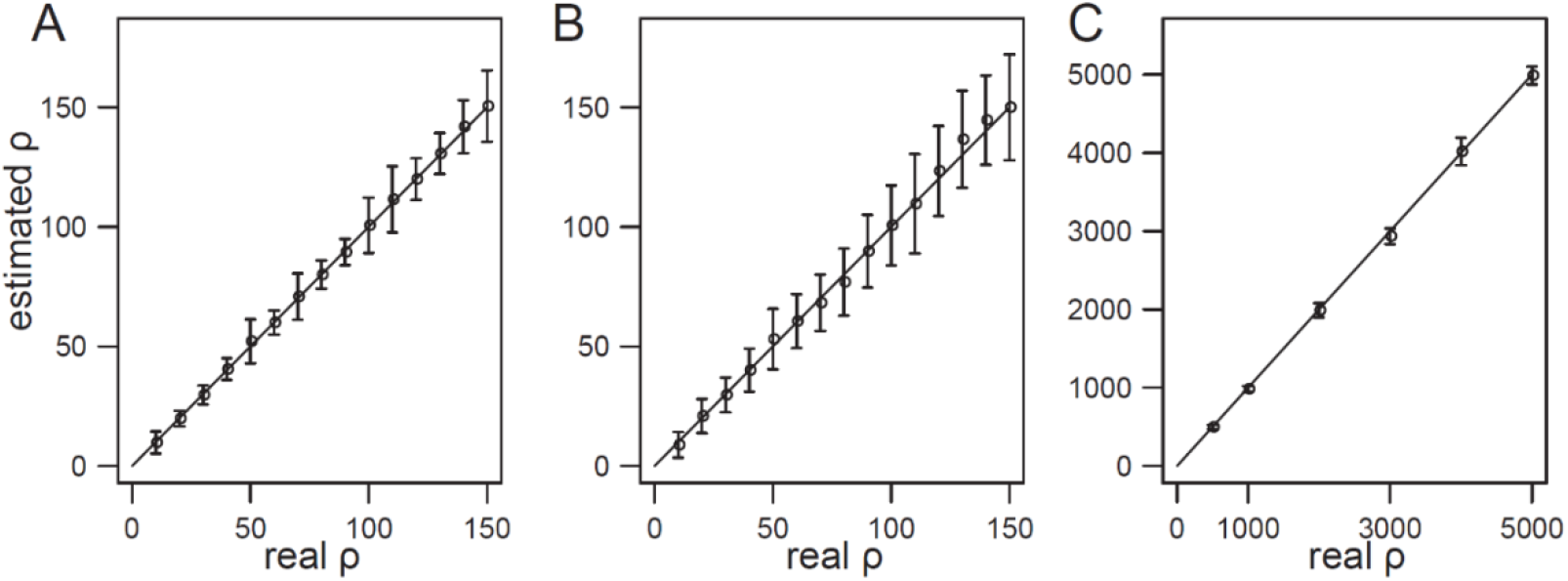
Comparison between estimated *ρ (ρ_FastEPRR_)* and real ρ under different conditions. (A) ρ ≤ 150, *n* = 8,000. (B) ρ ≤ 150, *n* = 200. (C) ρ ≥ 500, *n* = 8,000. The mean and the standard deviation of 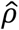 in (A) and (B) were calculated from 100 simulated data sets, and those in (C) were calculated from 10 simulated data sets. Error bars represent the standard deviation of 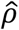.

We previously examined the cases 20 ≤ ρ ≤ 150 and found that the standard deviation of estimated 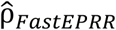 was smaller than that of 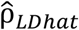^27^, suggesting that FastEPRR estimated the recombination rate more precisely than LDhat when recombination rate was not extremely low. However, it has been shown that certain chromosomal regions, such as recombination hotspots, have higher recombination rates than other regions. To assess the effectiveness of FastEPRR on these regions, the accuracy to infer high recombination rate was re-examined. Analysis of chromosomal regions with higher recombination rates (ρ ranging from 500 to 5,000) revealed that FastEPRR were precise and unbiased on potential recombination hotpot regions (Figure 1C).

FastEPRR provides the ability to infer recombination rate under different demographic models^27^, and it has been reported that demography affects estimated recombination rate^38–40^. Thus in this study we estimated the recombination rates of genomes in UK10K conditional on the Out-of-Africa model^32^. For the *i*-th chromosomal region, we need to estimate *N*_0_ first to calculate *r*_*i*_ from ρ_*i*_ (= 4*N*_0_*r*_*i*_). Therefore, the average ρ (= 4*N*_0_*r*) per Mb of 22 autosomes estimated by FastEPRR was 12,098.2, where *N*_0_ is the effective population size at present. As the average *r* of the same chromosomal regions in the 2019 deCODE family-based genetic map has been determined to be 1.2806 cM/Mb^18^, *N*_0_ was estimated as 235,186 for the UK10K data set under the Out-of-Africa model, in the range of estimated value in previous studies^32;41–43^. This value is 20.55 larger than the value previously estimated for the CEU population based on 1000 genomes in the OMNI CEU data set^44^ by LDhat^25^. This difference is reasonable because the current analysis was based on the Out-of-Africa model, whereas the previous one was based on the constant size model.

To compare the newly established genetic map with others, the Pairwise Pearson correlation coefficients between the UK10K map and 2019 deCODE family-based genetic map and the map of 1000 Genomes in the OMNI CEU data set were calculated at the 5-Mb and 1-Mb scales (Figure 2 and 3). The correlation coefficients were determined to be 0.8413 and 0.7256 at the 5-Mb scale, and 0.7308 and 0.5267 at the 1-Mb scale, indicating that the UK10K map is highly similar to the other two genetic maps.

**Figure 2.**
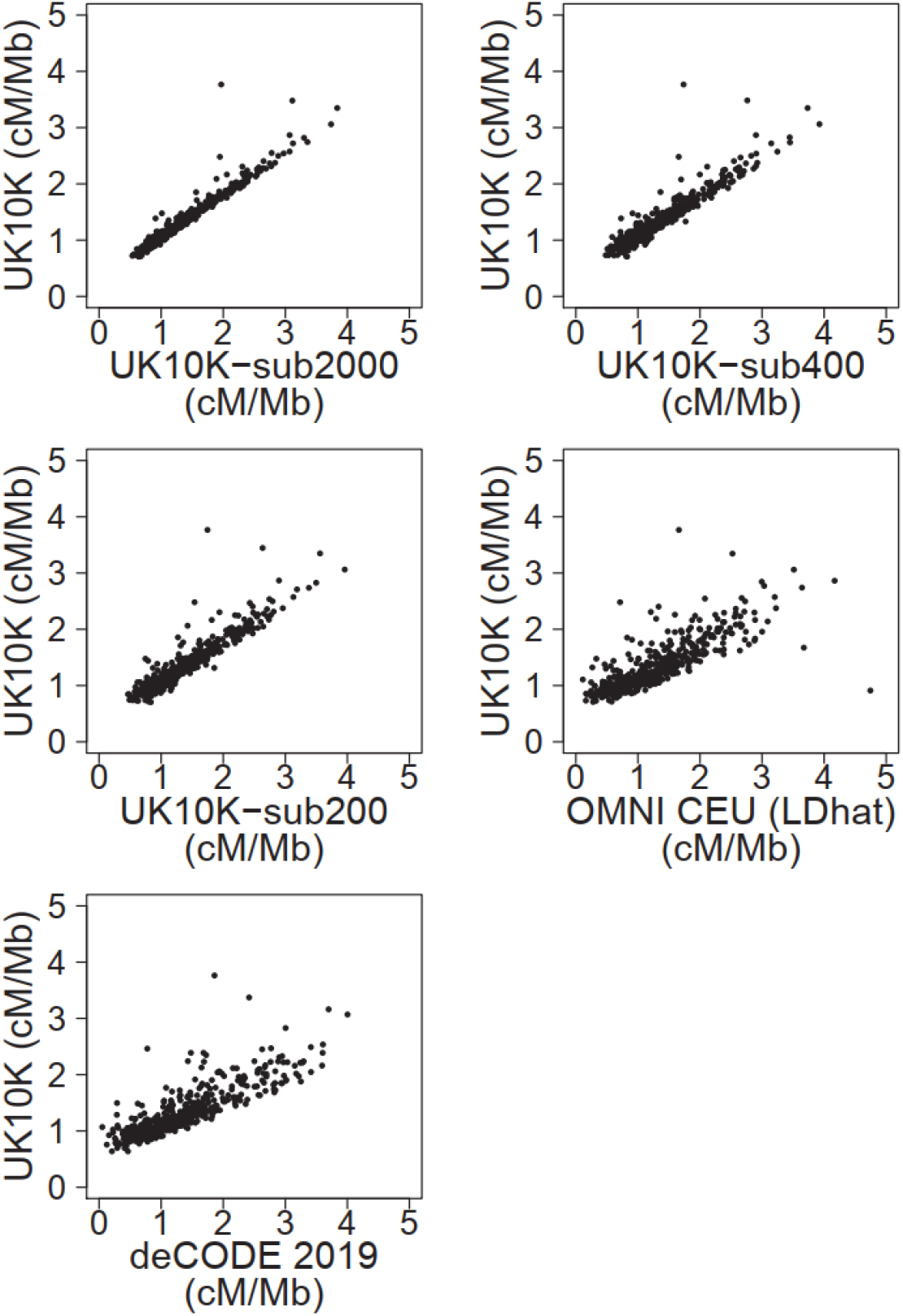
Correlation between the UK10K genetic map and five other maps at the 5-Mb scale. The recombination rates of UK10K, UK10K-sub2000, UK10K-sub400 and UK10K-sub200 were estimated by FastEPRR under the Out-of-Africa model, while those of OMNI CEU genomes were estimated previously by LDhat under the constant size model^44^. Each black dot represents two recombination rates of a window estimated by two different methods.

**Figure 3.**
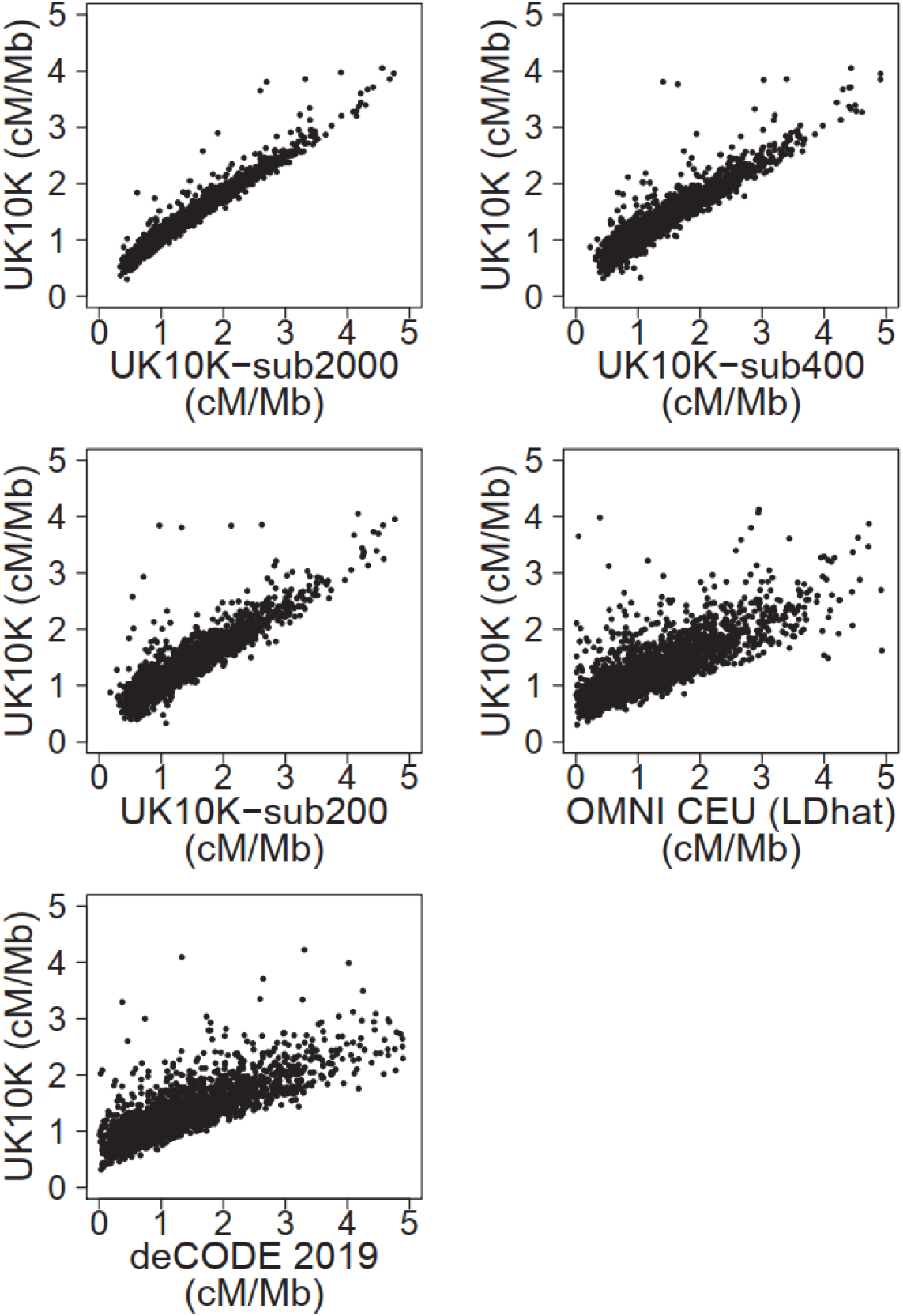
Correlation between the UK10K genetic map and five other maps at the 1-Mb scale. The recombination rates of UK10K, UK10K-sub2000, UK10K-sub400 and UK10K-sub200 were estimated by FastEPRR under the Out-of-Africa model, while those of OMNI CEU genomes were estimated previously by LDhat under the constant size model^44^. Each black dot represents two recombination rates of a window estimated by two different methods.

Genetic maps were also constructed under the Out-of-Africa model for the three subsets (UK10K-sub200, UK10K-sub400 and UK10K-sub2000) with 200, 400 and 2,000 genomes randomly selected from UK10K. The Pearson correlation coefficients between the UK10K and UK10K-sub200 maps were determined to be 0.9266 and 0.8216 at the 5- and 1-Mb scales, respectively, and those between the UK10K and UK10K-sub400 and UK10K-sub2000 maps were 0.9393 and 0.9658 at the 5-Mb scale, and 0.8501 and 0.9097 at the 1-Mb scale, respectively (Figure 2 and 3). These results indicate that the UK10K map and the maps of the three subsets are highly correlated with each other.

Recombination rates over different physical scales show that smaller scale has higher resolution (Figure 4), consistent with the previous finding^1^. To compare the distribution of recombination activities in different genetic maps, the recombination rate at the 5-kb scale was used. The recombination rates in chromosome 1 are shown as an example in Figure 5. Two regions (chr1:15,425,001-15,430,000 and chr1:236,680,001-236,685,000) were found to have higher recombination rates with seven PRDM9 (PR domain zinc finger protein 9) binding peaks^45^, indicative of the ability of FastEPRR to detect potential recombination hotspots. Since the human leukocyte antigen (HLA) region is one of the most polymorphic and recombined regions in human genome^46–49^, it was analyzed and multiple recombination hotspot candidates, hotter than the previous studies, were detected (Figure 6).

**Figure 4.**
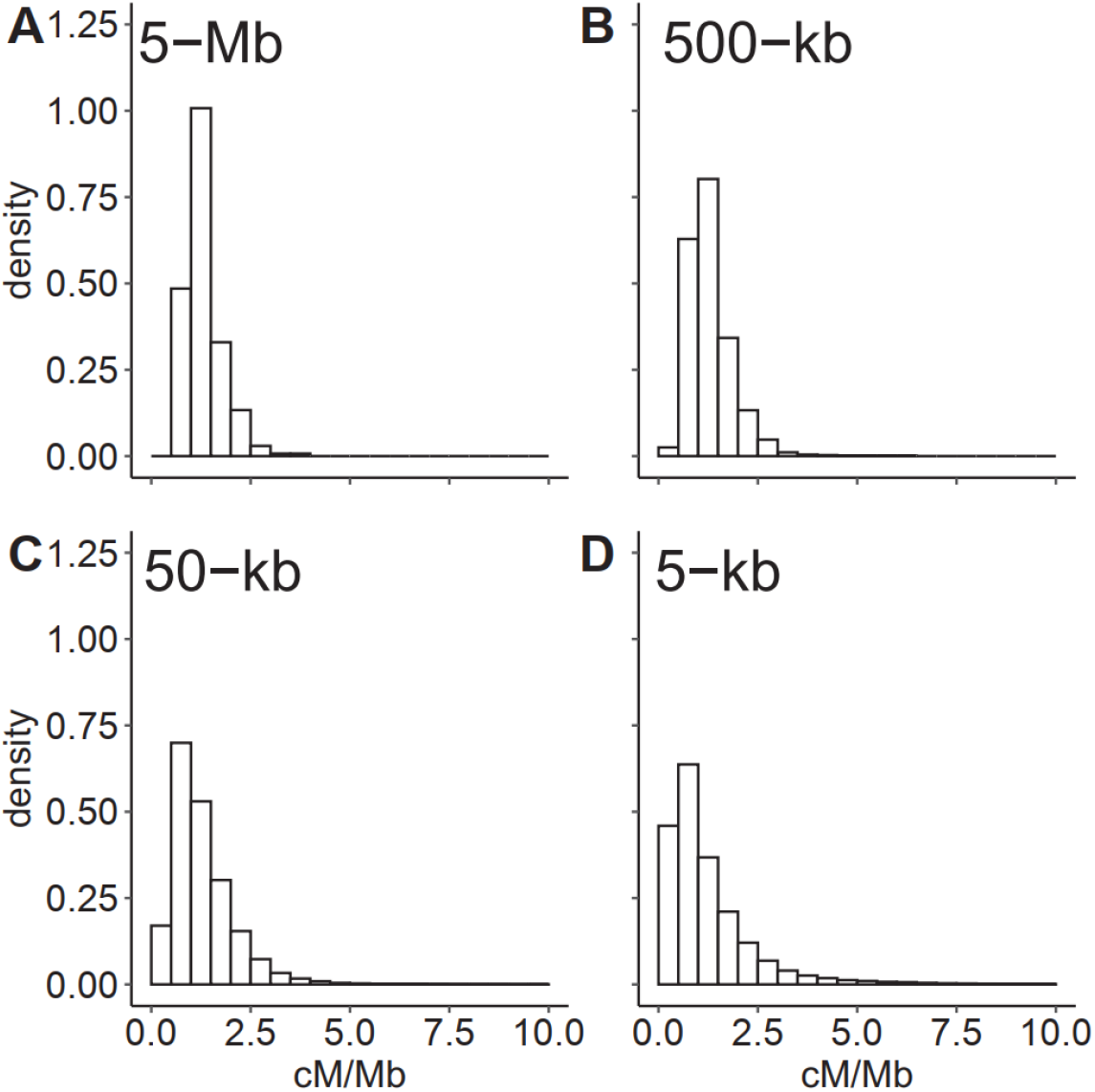
Recombination rates of the human genome at various physical scales. The skewed distribution indicates that the most of chromosomal regions have low recombination rate.

**Figure 5.**
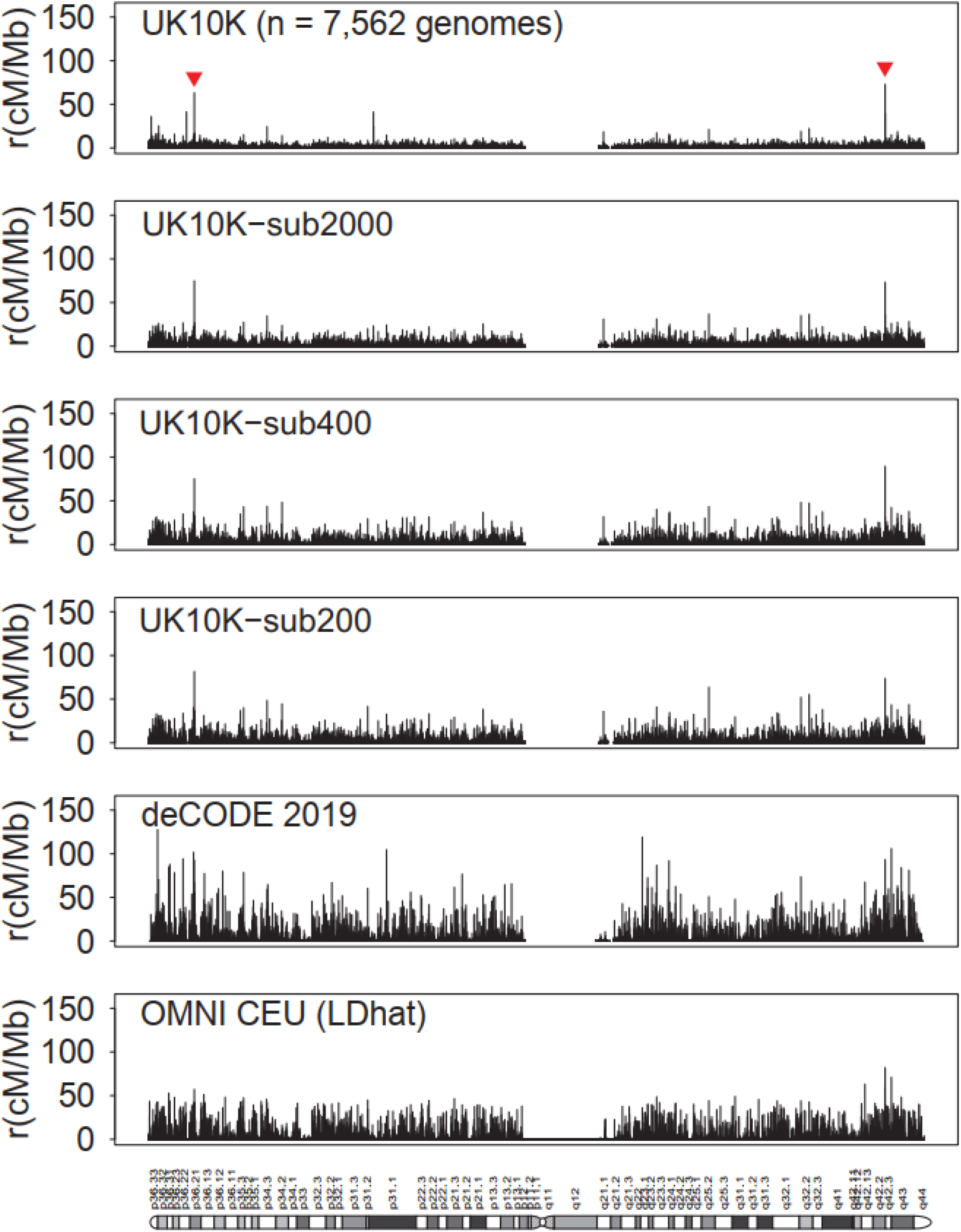
Recombination rates of various regions of human chromosome 1 obtained from six different genetic maps at the 5-kb scale. The height of each spike on the maps represents the average recombination rate of a 5-kb region in chromosome 1. The recombination rates of various regions in chromosome 1 shown in the maps of UK10K, UK10K-sub2000, UK10K-sub400 and UK10K-sub200 were estimated by FastEPRR with the Out-of-Africa model. The recombination rates of chromosome 1 in deCODE 2019 and the OMNI CEU data set estimated by LDhat^44^ with the constant size model are shown for comparison.

**Figure 6.**
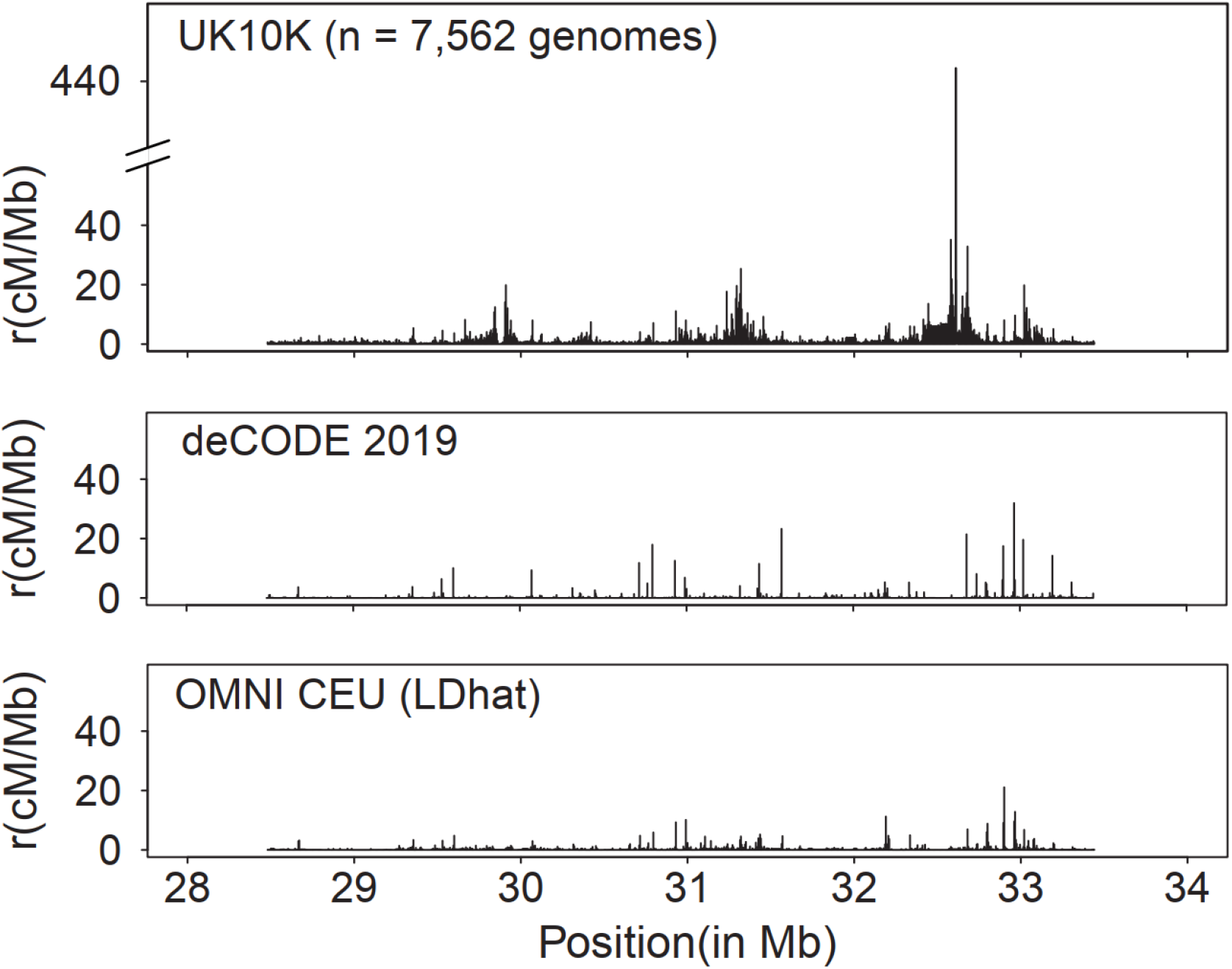
Recombination rates of various regions of the human HLA locus region estimated from three genetic maps at 5-kb scale. The height of each spike on the maps represents the average recombination rate of a 5-kb region in the HLA locus. The recombination rates of various regions shown in the maps of UK10K were estimated by FastEPRR with the Out-of-Africa model. The recombination rates of deCODE 2019 and the OMNI CEU data set estimated by LDhat^44^ with the constant size model are shown for comparison.

Although recombination hotspots candidates are revealed in the UK10K genetic map, the distribution of recombination events is generally less zigzag than other human genetic maps (Figure 5). The variance of recombination rate was then determined and found to be 3.4477 cM^2^/Mb^2^ in UK10K, 4.7224 cM^2^/Mb^2^ in UK10K-sub2000, 7.3097 cM^2^/Mb^2^ in UK10K-sub400, 7.9453 cM^2^/Mb^2^ in UK10K-sub200, 17.9188 cM^2^/Mb^2^ in OMNI CEU LDhat, and 25.7734 cM^2^/Mb^2^ in deCODE (calculated at the 5-kb scale). The variance of recombination rate in the UK10K genomes is the smallest. The proportion of total recombination in various percentages of the human genome was plotted and analyzed (Figure 7A). Overall, recombination activity in the UK10K genomes is much less concentrated than those of the genomes in other data sets. When the sample size was smaller than or equal to 400 chromosomes, 54.88-82.59% of crossover events were observed in 10% of the human genome. This observation is consistent with previous findings^18;27;50^. Only 33.82% of crossover events were detected in 10% of the human genome in UK10K. This percentage is strikingly lower than that of previous studies^18;27;50^. This difference is likely due to the more precise estimation of recombination rates with an extraordinarily large sample size.

**Figure 7.**
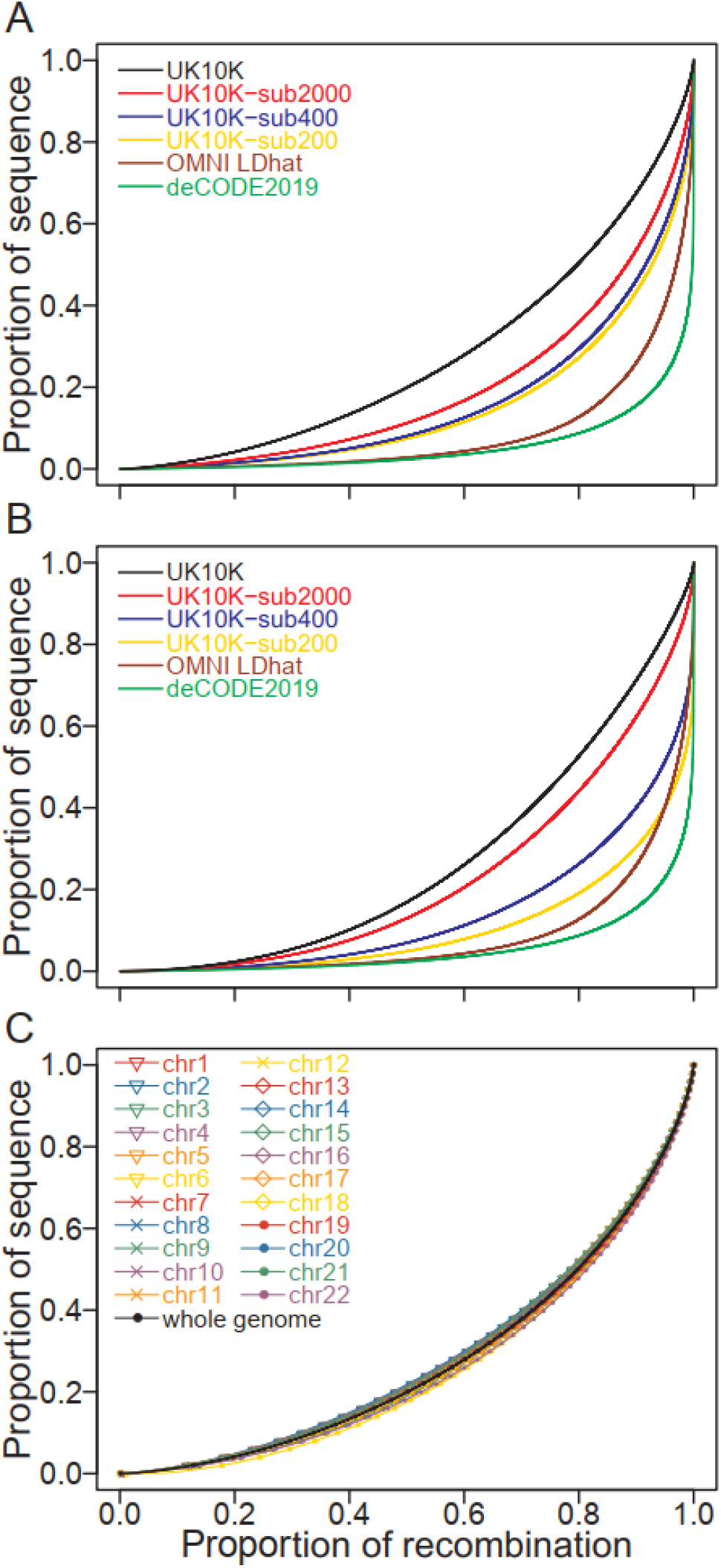
Proportion of recombination in different fractions of the sequence. (A) Recombination rates of genomes in various datasets estimated using the Out-of-Africa model. (B) Recombination rates of genomes in various datasets estimated using the constant size model. (C) Proportion of recombination in different fractions of the sequence of 22 autosomes compared to that of the whole genome. The recombination rates of various chromosomes were estimated using the Out-of-Africa model.

To valid our conclusions, the recombination rates were also estimated conditional on the constant size model. The Pearson correlation coefficient between the estimated recombination rate under the Out-of-Africa model and that under the constant model was only 0.7893 at the 5-kb scale, indicating that demography affects the estimation of recombination rates^27;38–40^. However, the newly estimated recombination rates are still less variable with larger sample size (Figure 7B), indicating that our conclusion is robust with the assumption of demography.

The reason why large sample yielded more accurate results was investigated. As more recombination events and mutations (equivalent to SNPs) occur on a tree with a longer tree length, both factors may affect the determination of recombination rates. To determine the effect of SNP numbers, two cases were investigated. In the first case (Case01), 200 genomes were selected from the samples in UK10K-sub400. In the second case (Case02), all 400 genomes in UK10K-sub400 were considered, however, only the SNPs in the Case01 were analyzed. In other words, these two cases had the same number of SNPs but had different sample sizes as well as different recombination events. Results showed that the estimated recombination rate of Case02 is slightly more similar to that of UK10K-sub400 than that of the Case01 (Correlation coefficient 0.9770 *vs* 0.9609 at the 5-kb scale). The variance of recombination rate of Case02 is also slightly smaller than that of Case01 (7.2850 cM^2^/Mb^2^ *vs* 7.5130 cM^2^/Mb^2^, calculated at the 5-kb scale). Similar results were obtained with the constant size model (Correlation coefficient 0.9692 *vs* 0.9562, and variance 10.5054 cM^2^/Mb^2^ *vs* 14.9622 cM^2^/Mb^2^, calculated at 5-kb scale). Therefore, more recombination events observed in data set with large sample size, rather than more SNPs, result in a more precise estimate of recombination rate.

## Discussion

In this study, we employed the software FastEPRR to estimate the recombination rate of various regions of the human genome using datasets with different sample sizes. These datasets included UK10K, UK10K-sub2000, UK10K-sub400 and UK10K-sub200 with 7,562, 2,000, 400, and 200 genomes, respectively. The mean recombination rate in these four datasets is determined to be the same but their standard deviation increases when the sample size is reduced. This result showed that the larger the sample size for the analysis, the smaller fluctuation of estimated recombination rate, and the less concentrated recombination activities of the human genome.

These findings are robust with the assumption of demographic models. Under the constant population size and the Out-of-Africa models, the same conclusions were made. These findings are also consistent with the expectation of coalescent theory, which indicates that the coalescent tree length of a larger sample is longer than that of a smaller sample. The genomes of the former thus contain more recombination events than those of the latter, leading a more precise estimate of recombination rate. In our study, the sample size is not very small comparing with the long-term human effective population size (~10,000). Although Hudson’s *ms* simulator requires that the sample size is much smaller than the effective population size^37^, our findings are robust with this assumption. For the exact coalescent^51^ and the Moran model^52^, given a true tree with the sample size (*n* − 1), the tree with the sample *n* can be obtained by adding a tip with a non-zero length, and thus the tree length increases. The same conclusion can be made under different demographic histories. Since FastEPRR is currently the only method which is fast enough to handle a super-large dataset, we cannot confirm the findings by using alternative methods. However, it has been shown that when the sample size was increased from 50 to 200 ^27;28^, the variance of recombination rate estimated by LDhat^25^, summary statistic likelihood (SSL)^53^ and product of approximate conditionals (PAC)^54^ was reduced. As our results generated by FastEPRR resemble those of these alternative methods, it is conceivable our findings represent a general phenomenon in humans and other sexually reproduced species.

This study is the first attempt to estimate recombination rate using an extraordinarily large population genetic data set. By processing genomes with a sample size of four orders of magnitude, we demonstrated that artificial intelligence approaches^28;55–57^ are effective for population genetic analysis. The results suggest that the fluctuation of recombination rate of human genome may be overestimated in previous studies.

## Supporting information

UK10K genetic map

## Data and Code Availability

The datasets used in this study are available at the UK10K Project Consortium^31^. The genetic maps of OMNI data set built by LDhat^25^ were downloaded from the 1000 Genomes Project. FastEPRR 2.0 is written in R and integrated on the eGPS cloud^58^ (http://www.egps-software.net). The desktop version and the genetic maps established in this study are freely available on the institute website (https://www.picb.ac.cn/evolgen/). The genetic maps are also available as supplemental materials.

## Web Resources

FastEPRR 2.0 and UK10K genetic map, https://www.picb.ac.cn/evolgen/softwares/UK10K Project Consortium, https://www.uk10k.org/

Genetic map of OMNI data set, ftp://ftp.1000genomes.ebi.ac.uk/vol1/ftp/technical/working/20130507_omni_recombination_rates, eGPS software, http://www.egps-software.net

## Acknowledgments

We thank the UK10K Project Consortium for sharing the data. This work was supported by grants from the National Natural Science Foundation of China (nos. 31100273, 31172073, 91131010), the Strategic Priority Research Program of the Chinese Academy of Sciences (Grant No. XDB38030100), and National Key Research and Development Project (No. 2020YFC0847000).

## References

1. Coop, G., and Przeworski, M. (2007). An evolutionary view of human recombination. Nat Rev Genet 8, 23–34.

2. Hill, W.G., and Robertson, A. (1968). Linkage disequilibrium in finite populations. Theoretical and Applied Genetics 38, 226–231.

3. Ohta, T., and Kimura, M. (1971). Linkage disequilibrium between two segregating nucleotide sites under the steady flux of mutations in a finite population. Genetics 68, 571–580.

4. Ardlie, K.G., Kruglyak, L., and Seielstad, M. (2002). Patterns of linkage disequilibrium in the human genome. Nat Rev Genet 3, 299–309.

5. Francioli, L.C., Polak, P.P., Koren, A., Menelaou, A., Chun, S., Renkens, I., van Duijn, C.M., Swertz, M., Wijmenga, C., van Ommen, G., et al. (2015). Genome-wide patterns and properties of de novo mutations in humans. Nat Genet 47, 822–826.

6. Kong, A., Thorleifsson, G., Frigge, M.L., Masson, G., Gudbjartsson, D.F., Villemoes, R., Magnusdottir, E., Olafsdottir, S.B., Thorsteinsdottir, U., and Stefansson, K. (2014). Common and low-frequency variants associated with genome-wide recombination rate. Nat Genet 46, 11–16.

7. Payseur, B.A., and Rieseberg, L.H. (2016). A genomic perspective on hybridization and speciation. Mol Ecol 25, 2337–2360.

8. Keinan, A., and Reich, D. (2010). Human population differentiation is strongly correlated with local recombination rate. Plos Genet 6, e1000886.

9. Price, A.L., Tandon, A., Patterson, N., Barnes, K.C., Rafaels, N., Ruczinski, I., Beaty, T.H., Mathias, R., Reich, D., and Myers, S. (2009). Sensitive detection of chromosomal segments of distinct ancestry in admixed populations. Plos Genet 5, e1000519.

10. Wegmann, D., Kessner, D.E., Veeramah, K.R., Mathias, R.A., Nicolae, D.L., Yanek, L.R., Sun, Y.V., Torgerson, D.G., Rafaels, N., Mosley, T., et al. (2011). Recombination rates in admixed individuals identified by ancestry-based inference. Nat Genet 43, 847–853.

11. Schumer, M., Xu, C.L., Powell, D.L., Durvasula, A., Skov, L., Holland, C., Blazier, J.C., Sankararaman, S., Andolfatto, P., Rosenthal, G.G., et al. (2018). Natural selection interacts with recombination to shape the evolution of hybrid genomes. Science 360, 656–659.

12. Stapley, J., Feulner, P.G.D., Johnston, S.E., Santure, A.W., and Smadja, C.M. (2017). Recombination: the good, the bad and the variable. Philos T R Soc B 372, 20170279.

13. Hernandez, R.D., Kelley, J.L., Elyashiv, E., Melton, S.C., Auton, A., McVean, G., Sella, G., Przeworski, M., and Project, G. (2011). Classic selective sweeps were rare in recent human evolution. Science 331, 920–924.

14. Stevison, L.S., Woerner, A.E., Kidd, J.M., Kelley, J.L., Veeramah, K.R., McManus, K.F., Bustamante, C.D., Hammer, M.F., Wall, J.D., and Project, G.A.G. (2016). The time scale of recombination rate evolution in Great Apes. Molecular Biology and Evolution 33, 928–945.

15. Hussin, J.G., Hodgkinson, A., Idaghdour, Y., Grenier, J.C., Goulet, J.P., Gbeha, E., Hip-Ki, E., and Awadalla, P. (2015). Recombination affects accumulation of damaging and disease-associated mutations in human populations. Nat Genet 47, 400–404.

16. Weiss, K.M., and Clark, A.G. (2002). Linkage disequilibrium and the mapping of complex human traits. Trends Genet 18, 19–24.

17. Kong, A., Thorleifsson, G., Gudbjartsson, D.F., Masson, G., Sigurdsson, A., Jonasdottir, A., Walters, G.B., Jonasdottir, A., Gylfason, A., Kristinsson, K.T., et al. (2010). Fine-scale recombination rate differences between sexes, populations and individuals. Nature 467, 1099–1103.

18. Halldorsson, B.V., Palsson, G., Stefansson, O.A., Jonsson, H., Hardarson, M.T., Eggertsson, H.P., Gunnarsson, B., Oddsson, A., Halldorsson, G.H., Zink, F., et al. (2019). Characterizing mutagenic effects of recombination through a sequence-level genetic map. Science 363, eaau1043.

19. Webb, A.J., Berg, I.L., and Jeffreys, A. (2008). Sperm cross-over activity in regions of the human genome showing extreme breakdown of marker association. P Natl Acad Sci USA 105, 10471–10476.

20. Arbeithuber, B., Betancourt, A.J., Ebner, T., and Tiemann-Boege, I. (2015). Crossovers are associated with mutation and biased gene conversion at recombination hotspots. P Natl Acad Sci USA 112, 2109–2114.

21. Bell, A.D., Mello, C.J., Nemesh, J., Brumbaugh, S.A., Wysoker, A., and McCarroll, S.A. (2020). Insights into variation in meiosis from 31,228 human sperm genomes. Nature 583, 259–264.

22. Hudson, R.R. (2001). Two-locus sampling distributions and their application. Genetics 159, 1805–1817.

23. Fearnhead, P., and Donnelly, P. (2001). Estimating recombination rates from population genetic data. Genetics 159, 1299–1318.

24. McVean, G., Awadalla, P., and Fearnhead, P. (2002). A coalescent-based method for detecting and estimating recombination from gene sequences. Genetics 160, 1231–1241.

25. Auton, A., and McVean, G. (2007). Recombination rate estimation in the presence of hotspots. Genome Res 17, 1219–1227.

26. McVean, G.A.T., Myers, S.R., Hunt, S., Deloukas, P., Bentley, D.R., and Donnelly, P. (2004). The fine-scale structure of recombination rate variation in the human genome. Science 304, 581–584.

27. Gao, F., Ming, C., Hu, W.J., and Li, H.P. (2016). New software for the fast estimation of population recombination rates (FastEPRR) in the genomic era. G3-Genes Genomes Genet. 6, 1563–1571.

28. Lin, K., Futschik, A., and Li, H.P. (2013). A fast estimate for the population recombination rate based on regression. Genetics 194, 473–484.

29. Sall, T., and Nilsson, N.O. (1994). The robustness of recombination frequency estimates in intercrosses with dominant markers. Genetics 137, 589–596.

30. Gartner, K., and Futschik, A. (2016). Improved versions of common estimators of the recombination rate. Journal of Computational Biology 23, 756–768.

31. Walter, K., Min, J.L., Huang, J., Crooks, L., Memari, Y., McCarthy, S., Perry, J.R.B., Xu, C., Futema, M., Lawson, D., et al. (2015). The UK10K project identifies rare variants in health and disease. Nature 526, 82–89.

32. Gravel, S., Henn, B.M., Gutenkunst, R.N., Indap, A.R., Marth, G.T., Clark, A.G., Yu, F.L., Gibbs, R.A., Bustamante, C.D., and Project, G. (2011). Demographic history and rare allele sharing among human populations. P Natl Acad Sci USA 108, 11983–11988.

33. Li, H.P., and Stephan, W. (2005). Maximum-likelihood methods for detecting recent positive selection and localizing the selected site in the genome. Genetics 171, 377–384.

34. Hudson, R.R. (2002). Generating samples under a Wright-Fisher neutral model of genetic variation. Bioinformatics 18, 337–338.

35. Hothorn, T., Buehlmann, P., Kneib, T., Schmid, M., and Hofner, B. (2018). mboost: Model-Based Boosting, R package version 2.9-1, https://CRAN.R-project.org/package=mboost.

36. R Core Team. (2019). R: A language and environment for statistical computing.

37. Kingman, J.F.C. (1982). On the genealogy of large populations. J Appl Probab 19, 27–43.

38. Spence, J.P., and Song, Y.S. (2019). Inference and analysis of population-specific fine-scale recombination maps across 26 diverse human populations. Science Advances 5, eaaw9206.

39. Kamm, J.A., Spence, J.P., Chan, J., and Song, Y.S. (2016). Two-locus likelihoods under variable population size and fine-scale recombination rate estimation. Genetics 203, 1381–1399.

40. Dapper, A.L., and Payseur, B.A. (2018). Effects of Demographic History on the Detection of Recombination Hotspots from Linkage Disequilibrium. Molecular Biology and Evolution 35, 335–353.

41. Terhorst, J., Kamm, J.A., and Song, Y.S. (2017). Robust and scalable inference of population history from hundreds of unphased whole genomes. Nat Genet 49, 303–309.

42. Schiffels, S., and Durbin, R. (2014). Inferring human population size and separation history from multiple genome sequences. Nat Genet 46, 919–925.

43. Hu, W.J., Hao, Z.Q., Du, P.Y., Di Vincenzo, F., Manzi, G., Pan, Y.H., and Li, H.P. (2021). Genomic inference of a human super bottleneck in Mid-Pleistocene transition. bioRxiv 444351.

44. Altshuler, D.M., Durbin, R.M., Abecasis, G.R., Bentley, D.R., Chakravarti, A., Clark, A.G., Donnelly, P., Eichler, E.E., Flicek, P., Gabriel, S.B., et al. (2012). An integrated map of genetic variation from 1,092 human genomes. Nature 491, 56–65.

45. Altemose, N., Noor, N., Bitoun, E., Tumian, A., Imbeault, M., Chapman, J.R., Aricescu, A.R., and Myers, S.R. (2017). A map of human PRDM9 binding provides evidence for novel behaviors of PRDM9 and other zinc-finger proteins in meiosis. eLife 6, e28383.

46. Wu, R.G., Li, H.X., Peng, D., Li, R., Zhang, Y.M., Hao, B., Huang, E.W., Zheng, C.H., and Sun, H.Y. (2019). Revisiting the potential power of human leukocyte antigen (HLA) genes on relationship testing by massively parallel sequencing-based HLA typing in an extended family. J Hum Genet 64, 29–38.

47. Jeffreys, A.J., Kauppi, L., and Neumann, R. (2001). Intensely punctate meiotic recombination in the class II region of the major histocompatibility complex. Nat Genet 29, 217–222.

48. Cullen, M., Perfetto, S.P., Klitz, W., Nelson, G., and Carrington, M. (2002). High-resolution patterns of meiotic recombination across the human major histocompatibility complex. Am J Hum Genet 71, 759–776.

49. Miretti, M.M., Walsh, E.C., Ke, X.Y., Delgado, M., Griffiths, M., Hunt, S., Morrison, J., Whittaker, P., Lander, E.S., Cardon, L.R., et al. (2005). A high-resolution linkage-disequilibrium map of the human major histocompatibility complex and first generation of tag single-nucleotide polymorphisms. Am J Hum Genet 76, 634–646.

50. Myers, S., Bottolo, L., Freeman, C., McVean, G., and Donnelly, P. (2005). A fine-scale map of recombination rates and hotspots across the human genome. Science 310, 321–324.

51. Fu, Y.X. (2006). Exact coalescent for the Wright-Fisher model. Theor Popul Biol 69, 385–394.

52. Wirtz, J., and Wiehe, T. (2019). The evolving Moran genealogy. Theor Popul Biol 130, 94–105.

53. Wall, J.D. (2000). A comparison of estimators of the population recombination rate. Molecular Biology and Evolution 17, 156–163.

54. Li, N., and Stephens, M. (2003). Modeling linkage disequilibrium and identifying recombination hotspots using single-nucleotide polymorphism data. Genetics 165, 2213–2233.

55. Lin, K., Li, H.P., Schlotterer, C., and Futschik, A. (2011). Distinguishing positive selection from neutral evolution: Boosting the performance of summary statistics. Genetics 187, 229–244.

56. Flagel, L., Brandvain, Y., and Schrider, D.R. (2019). The unreasonable effectiveness of convolutional neural networks in population genetic inference. Molecular Biology and Evolution 36, 220–238.

57. Pavlidis, P., Jensen, J.D., and Stephan, W. (2010). Searching for footprints of positive selection in whole-genome SNP data from nonequilibrium populations. Genetics 185, 907–922.

58. Yu, D.L., Dong, L.L., Yan, F.Q., Mu, H.L., Tang, B.X., Yang, X., Zeng, T., Zhou, Q., Gao, F., Wang, Z.H., et al. (2019). eGPS 1.0: Comprehensive software for multi-omic and evolutionary analyses. National Science Review 6, 867–869.

